# Chemokine Signaling Shapes Hepatic Lipid Homeostasis through the CXCL12/CXCR4/CXCR7 Axis

**DOI:** 10.64898/2025.12.03.692068

**Authors:** Rachel Barkan-Michaeli, Chaim Atay Fainshtein, Hadar Bar-Dagan, Nof Hadar, Einat Cinnamon, Ilan Stein, Kfir Sharabi

## Abstract

The CXCL12/CXCR4/CXCR7 signaling axis, long recognized for its roles in cancer, fibrosis, and tissue repair, is emerging as a broader regulator of tissue homeostasis. Here, we uncover a previously unappreciated function of this pathway in regulating hepatic metabolism. We show that *Cxcl12*, *Cxcr4*, and *Ackr3* (encoding CXCR7) are dynamically regulated during the fasting-refeeding transition and become dysregulated under conditions of diet-induced insulin resistance. Hepatocyte-specific depletion of *Cxcl12* or overexpression of *Ackr3* each led to hepatic triglyceride accumulation, whereas hepatocyte *Cxcr4* overexpression did not reproduce this phenotype, supporting a nonredundant role for CXCR7 in hepatocytes. Hepatic SDF-1α measurements across the three in vivo models further supported coordinated regulation of ligand availability within the axis, including a marked reduction in the Ackr3 overexpression model, consistent with enhanced ligand scavenging. In primary hepatocytes, *Ackr3* overexpression promoted lipid accumulation and was associated with altered AKT-linked signaling, particularly under lipid-rich conditions. Analysis of human liver transcriptomic datasets revealed reduced *CXCL12* and elevated *CXCR4* and *ACKR3* expression in NAFLD and NASH, supporting the translational relevance of this pathway. Together, our results identify the CXCL12/CXCR4/CXCR7 axis as an integral regulator of hepatic lipid balance and highlight CXCR7 as a potential therapeutic target for metabolic liver disease.

## Introduction

The liver is central in maintaining whole-body glucose and lipid homeostasis, functioning as a critical hub for metabolic regulation(1–3). It orchestrates glucose production through glycogenolysis and gluconeogenesis while also regulating lipid metabolism by balancing lipogenesis, fatty acid oxidation, and lipoprotein production. These tightly coordinated processes ensure energy availability during both fed and fasting states, thereby supporting overall metabolic equilibrium. Disruptions in hepatic metabolism, however, can lead to the pathogenesis of metabolic disorders, such as type 2 diabetes (T2D) and metabolic-associated steatotic liver disease (MASLD)(4). Excessive hepatic glucose production and impaired insulin sensitivity are hallmark features of T2D, a condition characterized by chronic hyperglycemia and systemic insulin resistance. Similarly, dysregulated lipid metabolism underlies MASLD, formerly known as non-alcoholic fatty liver disease (NAFLD), which encompasses conditions ranging from benign steatosis to Metabolic dysfunction-associated steatohepatitis (MASH) and fibrosis(4). These metabolic dysfunctions compromise liver health and exacerbate systemic metabolic derangements, underscoring the liver’s pivotal role in maintaining metabolic balance and influencing disease pathogenesis.

The SDF1/CXCR4/CXCR7 signaling axis is increasingly recognized for its diverse roles in cancer, encompassing a broad spectrum of malignancies(5–10). Stromal cell-derived factor 1 (SDF1, also known as CXCL12) binds to its two receptors, CXCR4 and CXCR7, initiating distinct yet interrelated signaling cascades that govern tumor cell survival, proliferation, migration, and metastasis(10). CXCL12 is often overexpressed in tumor-associated stromal cells, thereby establishing chemokine gradients that attract *Cxcr4*-expressing cancer cells and support a protumorigenic microenvironment (11). CXCR4, which primarily signals through canonical G-protein–coupled pathways, has been extensively implicated in organ-specific metastasis, directing the migration of tumor cells toward CXCL12-rich tissues, such as the bone, liver, and lung(12). In contrast, CXCR7, initially considered a decoy or scavenger receptor, has emerged as a multifunctional atypical GPCR that not only regulates extracellular CXCL12 availability but also activates β–arrestin–dependent signaling pathways that promote angiogenesis, cell survival, and tumor growth(13). Recent mechanistic studies have revealed that CXCR7 (also known as ACKR3) can “sense” CXCR4 activation through GRK-dependent phosphorylation events, enabling dynamic receptor crosstalk that fine-tunes chemokine scavenging and β-arrestin recruitment(14, 15). This kinase-mediated feedback mechanism allows CXCR7 to modulate CXCR4 activity, thereby influencing both chemokine gradient formation and downstream cellular responses(14, 15). Collectively, these findings highlight that the CXCL12/CXCR4/CXCR7 axis constitutes a tightly interconnected regulatory network whose coordinated activity contributes to tumor progression, metastasis, and therapy resistance.

In the liver, CXCL12 is secreted by hepatic stellate cells, liver sinusoidal endothelial cells (LSECs), and malignant hepatocytes, particularly in response to injury(16). The CXCL12/CXCR4/CXCR7 signaling axis exerts dual, context-dependent roles, promoting regeneration following acute injury while driving fibrosis and tumor progression under chronic conditions(17). Activation of CXCR7 in LSECs enhances regenerative, pro-angiogenic signaling that supports hepatocyte proliferation, whereas CXCR4 activation during prolonged injury promotes fibrogenesis by recruiting hepatic stellate cells and stimulating extracellular matrix deposition(17). This balance is further influenced by factors such as FGF-2-mediated suppression of CXCR7, which shifts signaling toward CXCR4-driven fibrotic remodeling. Additionally, CXCL12/CXCR4 signaling contributes to the progression of hepatocellular carcinoma (HCC) by enhancing tumor cell survival, angiogenesis, and metastasis(16). Through such coordinated and context-specific activity, CXCR4 and CXCR7 collectively shape hepatic responses to injury and malignancy, balancing regenerative, fibrotic, and oncogenic programs within the liver microenvironment. This signaling plasticity highlights the therapeutic potential of selectively targeting the CXCL12/CXCR4/CXCR7 axis, as interventions must consider the dynamic interplay between these pathways to promote regeneration without exacerbating fibrosis or tumor progression.

Emerging evidence suggests that the CXCL12/CXCR4/CXCR7 pathway also plays a significant role in modulating insulin signaling. CXCL12 acts as both a systemic and local regulator of glucose metabolism, with its effects varying by tissue context. In adipocytes, SDF-1 is an autocrine factor that desensitizes insulin signaling by inducing extracellular signal-regulated kinase (ERK)-mediated phosphorylation and degradation of IRS-1, thereby reducing insulin-stimulated glucose uptake(18). This autocrine regulation exacerbates insulin resistance under conditions of obesity and fasting(18). Interestingly, obesity increases CXCR7 expression in adipose tissue macrophages (ATMs), which drives SDF-1-induced chemotaxis and inflammation through NF-κB regulation. Neutralizing CXCR7 reduces ATM infiltration, thereby improving inflammation and insulin resistance in obesity(19). Conversely, in pancreatic β-cells, SDF-1 supports cell survival and enhances insulin secretion by activating the Akt pathway, highlighting its dualistic effects(20). In brown adipose tissue, CXCL12 enhances thermogenesis and glucose utilization through CXCR4-mediated activation of p38 and ERK signaling, underscoring its role in energy expenditure(21). In primary hepatocytes, SDF-1α and SDF-1β, the two major splice variants of SDF-1, increase Akt phosphorylation, suppress the expression of gluconeogenic genes, *Pck1* and *G6pc*, and reduce glucose secretion into the medium(22). These findings reveal a complex interplay where the SDF-1 axis influences insulin sensitivity and glucose homeostasis, depending on the metabolic and cellular environment. Further research is needed to leverage these insights to develop therapeutic strategies that improve metabolic health in individuals with insulin resistance.

While the CXCL12/CXCR4/CXCR7 signaling axis is well established in cancer biology and liver injury responses, its role in normal liver physiology, particularly in hepatocyte metabolic regulation, remains poorly understood. Here, we examined this pathway as an integrated chemokine signaling axis in hepatocytes rather than focusing on a single receptor in isolation. We show that *Cxcl12*, *Cxcr4*, and *Ackr3* are dynamically regulated during the fasting-refeeding transition and become dysregulated under conditions of diet-induced metabolic stress. Using hepatocyte-specific in vivo manipulations, we further demonstrate that reduced hepatocyte-derived CXCL12 and increased hepatocyte ACKR3 each promote hepatic triglyceride accumulation, whereas hepatocyte *Cxcr4* overexpression does not reproduce this phenotype. Mechanistically, our findings link CXCR7 to altered mTORC1 signaling and hepatocyte lipid storage. Together, these results identify a nutritionally regulated, hepatocyte-autonomous role for the CXCL12/CXCR4/CXCR7 axis in hepatic lipid homeostasis and distinguish CXCR7 as a nonredundant regulator within this pathway.

## Results

### The CXCL12/CXCR4/CXCR7 axis is regulated in the fast-feeding transition

The transition between fasting and refeeding is a complex physiological process primarily regulated by insulin and glucagon. Identifying genes and pathways involved in this transition has significantly enhanced our understanding of hepatic adaptation to metabolic changes(23). Nonetheless, our knowledge of how chemokine networks, and particularly the CXCL12/CXCR4/CXCR7 axis, are regulated in this context remains limited. We observed that glucagon robustly induces the expression of *Ackr3*, which encodes the atypical GPCR CXCR7, in primary mouse hepatocytes, whereas insulin does not affect its expression (Figure 1A). This observation aligns with a recent RNA-seq study, which identified *Ackr3* as the ninth most upregulated gene following glucagon stimulation and the only GPCR among the top 30 induced genes (24). To investigate this axis further, we examined the regulation of *Cxcr4*, encoding CXCR4. Glucagon did not affect *Cxcr4* expression, whereas insulin significantly reduced its expression (Figure 1B), consistent with findings in primary human endothelial cells(25).

**Figure 1.**
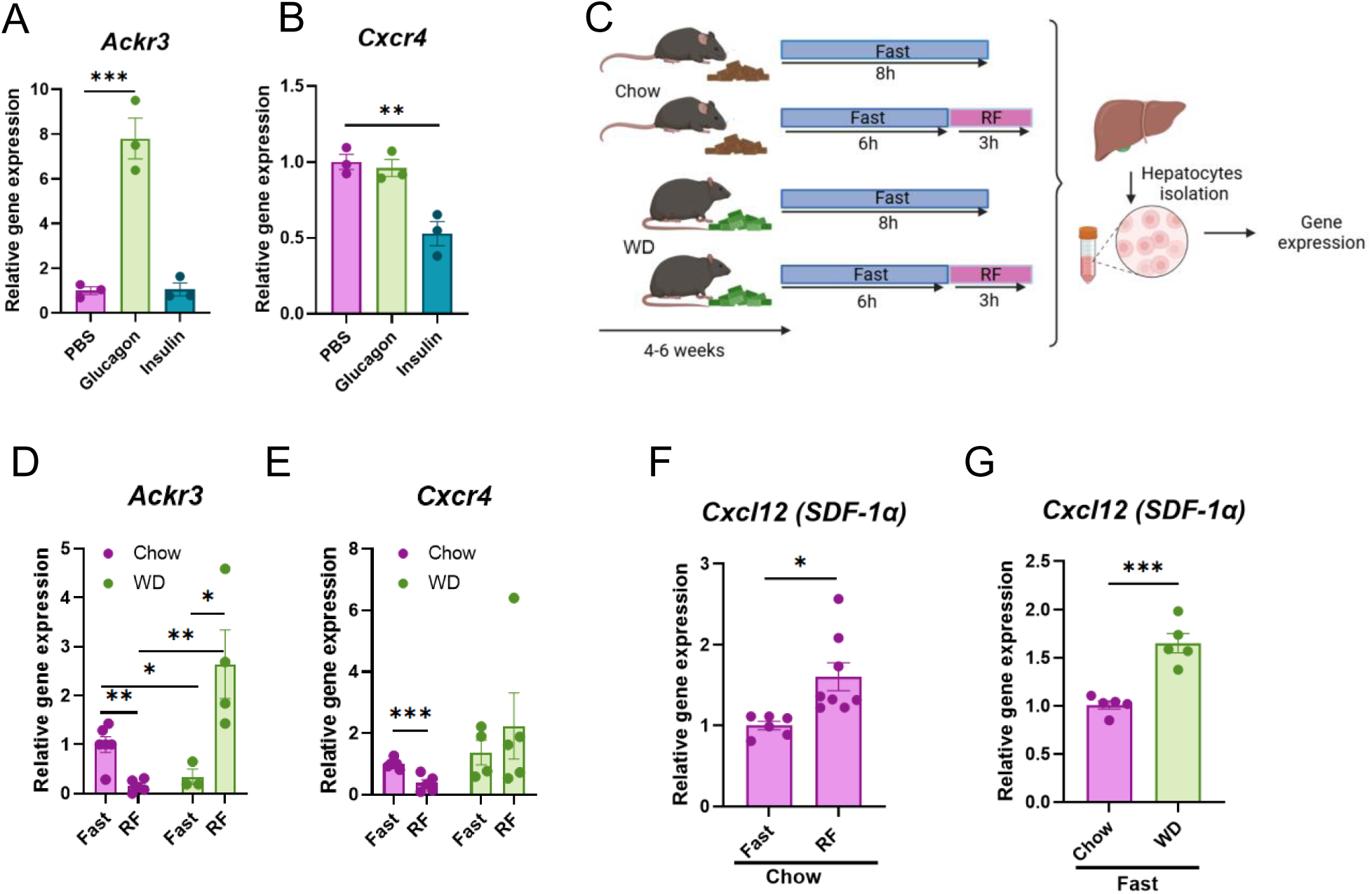
**(A)** Glucagon stimulates the expression of *Ackr3,* while **(B)** insulin suppresses the expression of *Cxcr4* in cultured mouse primary hepatocytes. **(C)** Fasting-refeeding experimental design: 6-week-old C57BL/6 mice were fed either chow or a Western diet. Within each experiment, all mice were exposed to the diet for the same duration and analyzed together, although the exact duration varied slightly between independent cohorts. Primary hepatocytes were isolated following an 8-hour fast, or a 6-hour fast followed by 3 hours of refeeding. (**D, E**) The expression level of *Ackr3* and *Cxcr4* in the hepatocytes fraction was measured. Results are a combination of 3 independent experiments with 1-3 mice in each group. **(F)** The expression level of *Cxcl12* from total liver extracts from chow-fed mice after an overnight fast, or an overnight fast followed by 3 hrs. refeeding. **(G)** The expression level of *Cxcl12* in total liver extracts following 6 hrs. fasting of chow-fed or WD-fed mice. *, p-value<0.05; **, p-value<0.01; ***, p-value<0.001.

In the liver, CXCR4 and CXCR7 are predominantly expressed in non-parenchymal cells (NPCs)(16). Thus, total liver lysate analyses mainly reflect NPC expression. To determine if the observed effects in cultured hepatocytes also occur *in vivo*, we examined the fasting-refeeding transition in adult C57BL/6 mice, with a specific focus on hepatocytes. Following 6 hours of fasting and 3 hours of refeeding, hepatocytes were separated from the NPCs, using collagenase digestion, and analyzed for gene expression (Figure 1C). Consistent with the *in vitro* findings, *Ackr3* mRNA expression levels in hepatocytes were significantly higher in the fasted state compared to refeeding (Figure 1D), suggesting that elevated *Ackr3* expression parallels elevated glucagon (fasting). At the same time, *Cxcr4* mRNA expression levels were significantly lower in hepatocytes in the refed state (Figure 1E), suggesting that decreased *Cxcr4* expression parallels elevated insulin (refeeding).

To study this axis under conditions that promote metabolic stress, we placed mice on a high-fat, high-sucrose Western diet (WD) for 4–6 weeks, a regimen commonly used to induce insulin resistance(26). *Ackr3* and *Cxcr4* expression were similarly assessed during fasting and refeeding to determine their expression specifically in hepatocytes. In WD-fed mice, fasting-induced *Ackr3* expression was significantly lower than in chow-fed controls. However, refeeding markedly increased *Ackr3* expression, exceeding fasting levels in chow-fed mice (Figure 1D). Conversely, *Cxcr4* expression during fasting was similar between WD-fed and chow-fed mice. Unlike chow-fed mice, *Cxcr4* expression in WD-fed mice remained unchanged during refeeding (Figure 1E). We also analyzed *Cxcl12* (encoding SDF-1), the primary ligand for CXCR7 and CXCR4. While NPCs, including HSCs and LSECs, are recognized as prominent producers of SDF-1α, hepatocytes also make a meaningful contribution to its hepatic pool (16, 27). To account for these contributions, we measured *Cxcl12* expression in whole liver tissue, focusing on the SDF-1α variant, the predominant form in the liver (28). Under chow diet conditions, *Cxcl12* (*SDF-1α*) expression was elevated during refeeding (Figure 1F). Interestingly, in WD-fed mice, *Cxcl12* (*SDF-1α*) expression was significantly higher during fasting compared to chow-fed mice (Figure 1G).

These results demonstrate that *Ackr3* and *Cxcr4* are regulated in hepatocytes *in vivo*, in a manner that parallels the opposing hormonal milieu of fasting and refeeding, and that their expression is further perturbed under metabolic stress induced by WD feeding. Additionally, *Cxcl12* is subject to transcriptional regulation during fasting and refeeding, as well as in insulin-resistant states. These findings highlight the potential significance of the CXCL12/CXCR4/CXCR7 axis in liver metabolism, underscoring the need for further investigation into its role in hepatic function and adaptation.

### *Cxcl12* downregulation promotes fat accumulation in WD-fed mice

To explore the role of the CXCL12/CXCR4/CXCR7 axis in hepatic metabolic adaptation, we assessed the effects of lowering *Cxcl12* expression levels. We utilized *Cxcl12*-flox mice to delete *Cxcl12* specifically in hepatocytes by injecting AAV-*Tbg*::Cre via the tail vein (Figure 2A). Although the NPCs are considered a significant source of CXCL12, *Cxcl12* is also expressed in healthy hepatocytes (16, 27). Deletion of *Cxcl12* in hepatocytes resulted in a substantial (∼60%) but not complete reduction of its expression in total liver lysates. The residual *Cxcl12* expression most likely originates from NPCs, as its expression in this fraction was only marginally affected by Cre recombinase activity driven by the hepatocyte-specific TBG promoter (Figures 2B and S1). This reduction in *Cxcl12* expression slightly but significantly increased *Cxcr4* expression without altering *Ackr3* expression (Figure 2C). Consistent with the reduction in hepatic *Cxcl12* expression, hepatic SDF-1α levels were also decreased in *Cxcl12*-deficient livers (Figure 2D).

**Figure 2.**
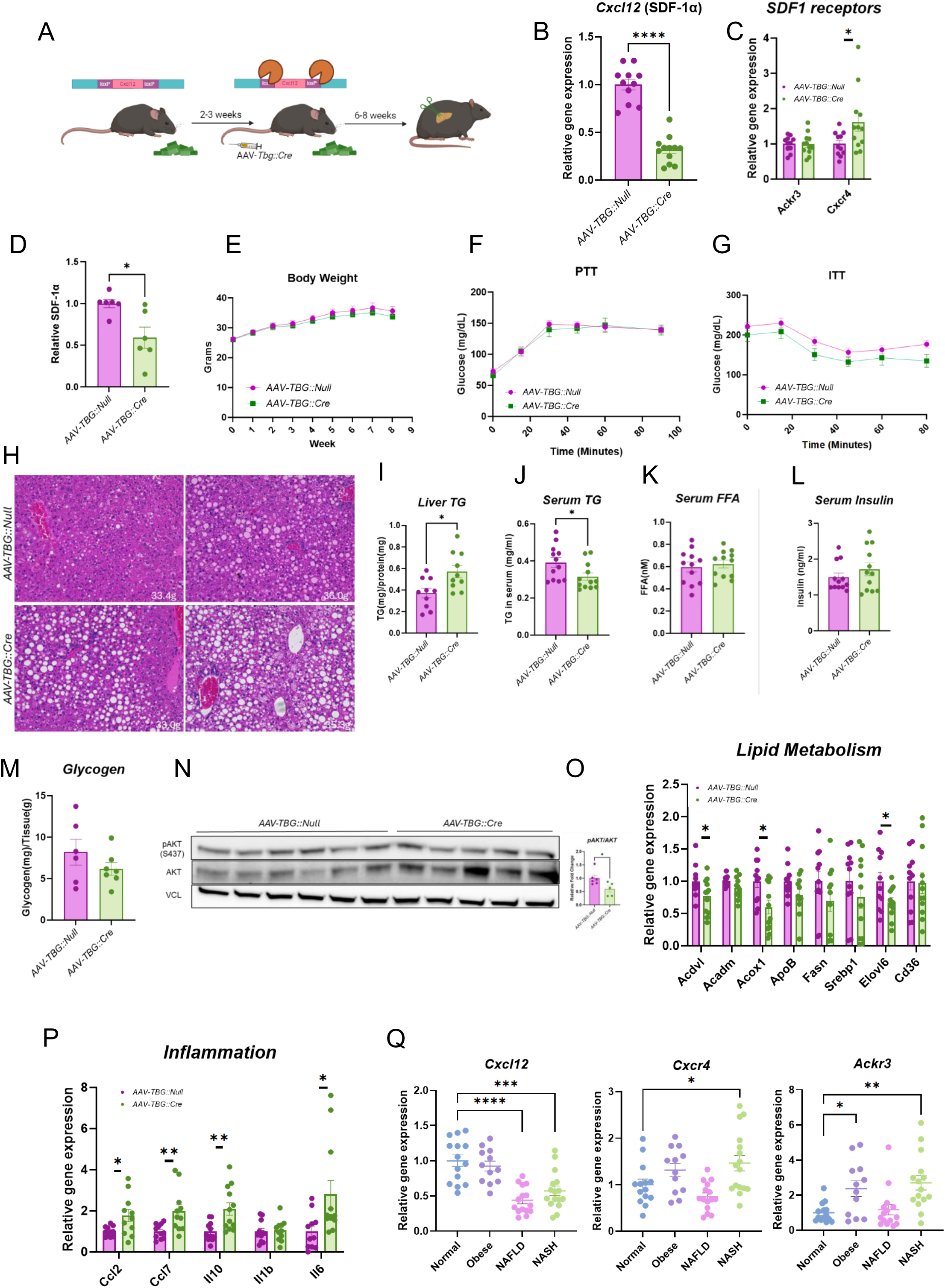
(**A**) Experimental design. *Cxcl12*-flox mice were placed on a Western diet beginning at 6 weeks of age. Within each experiment, all mice were exposed to the diet for the same duration and analyzed together, although the exact duration varied slightly between independent cohorts. Mice were injected with AAV-TBG::Null or AAV-TBG::Cre as indicated. Liver and serum samples were collected after 6hrs. of fasting. (**B**) *Cxcl12* expression level in the liver of control (*AAV-TBG::Null*) and hepatocytes-specific KO of *Cxcl12* (*AAV-TBG::Cre*) mice. (**C**) *SDF1* receptors expression level in the liver. (**D**) SDF-1α levels in total liver lysates following SDF-1α ELISA. (**E**) Body weight. (**F, G**) Pyruvate and insulin tolerance tests. (**H**) H&E staining of liver sections from *Cxcl12*-depleted mice. (**I**) Quantification of liver TG (**J**), serum triglycerides (**K**), serum FFA (**L**), and serum Insulin. (**M**) Liver glycogen content. (**N**) pAKT (S473) in total liver lysates. (**O**) Expression level of lipid metabolism-related and (**P**) inflammation-related genes. (**Q**) Expression level of *Cxcl12/Cxcr4/Ackr3* from human liver biopsies (Public database (GSE126848) by GEO2R online tool). *, p-value<0.05; **, p-value<0.01, ****, p-value<0.0001.

To assess the metabolic consequences of reduced hepatic CXCL12 levels, mice were fed a WD starting at six weeks of age for 2–3 weeks, followed by tail vein injection of AAV-*Tbg*::Cre or AAV-*Tbg*::null as a control (Figure 2A). Body weight gain was comparable between groups (Figure 2E), and no significant differences were observed in pyruvate tolerance tests (PTT) or insulin tolerance tests (ITT) (Figure 2F, G). However, histological analysis of H&E-stained liver sections revealed a marked increase in the size and number of lipid droplets in the livers of *Cxcl12**-***downregulated mice compared to controls (Figure 2H). Consistently, liver triglyceride (TG) quantification confirmed elevated TG levels in the knockout group (Figure 2I), while serum TG levels were decreased (Figure 2J), suggesting altered lipid metabolism. Serum free-fatty-acid (FFA) levels (Figure 2K), serum insulin (Figure 2L), and hepatic glycogen content (Figure 2M) were not significantly altered. Basal hepatic AKT phosphorylation following 6 hours of fasting was also reduced in livers from *Cxcl12*-deficient mice (Figure 2N), consistent with an effect of reduced ligand availability on downstream AKT signaling in vivo. Similar results regarding the accumulation of TG were also observed following depletion of *Cxcl12* using shRNA (Figure S2).

Gene expression analysis of lipid metabolism-related pathways revealed downregulation of *Acox1* and *Acadvl*, key genes involved in β-oxidation, in livers with reduced *Cxcl12* expression (Figure 2O). This downregulation likely contributes to hepatic fat accumulation. Additionally, *Elovl6*, a gene involved in fatty acid elongation, was also downregulated. Expression analysis revealed changes in a subset of inflammatory markers, including increased *Ccl2*, *Ccl7*, *Il10*, and *Il6* expression in livers with reduced *Cxcl12* (Figure 2P). However, these transcriptional changes were not accompanied by clear evidence of increased hepatic lymphocyte accumulation, as CD19 and CD3 staining were comparable between groups (Figure S3). In addition, F4/80 staining appeared reduced in *Cxcl12*-depleted livers (Figure S3). Together, these findings do not support robust inflammatory cell infiltration, but rather suggest a modest and heterogeneous inflammatory response associated with *Cxcl12* downregulation. Finally, analysis of publicly available human liver biopsy data (GSE126848, using the GEO2R online tool) indicates that the CXCL12/CXCR4/CXCR7 axis is also altered in human liver disease. In this dataset, which includes samples from individuals with obesity, NAFLD, and NASH, *Cxcl12* expression was reduced in NAFLD and NASH, while *Cxcr4* and *Ackr3* were elevated in NASH (Figure 2Q). These shifts resemble the patterns observed in our experimental models, suggesting that dysregulation of this chemokine axis may be a feature of human steatohepatitis. Taken together, these findings suggest that alterations in the CXCL12/CXCR4/CXCR7 axis may contribute to the inflammatory and metabolic remodeling characteristic of advanced liver disease, and highlight this pathway as a potential marker of disease progression.

Together, these data show that hepatocyte-derived *Cxcl12* influences hepatic lipid accumulation and inflammation, with similar alterations in the CXCL12/CXCR4/CXCR7 axis observed in human NAFLD and NASH.

### Hepatocyte-specific overexpression of *Ackr3* modulates hepatic lipid and glucose metabolism in WD-fed mice

The increase in liver fat content following reduced expression of *Cxcl12* may result from changes in the activity of its downstream receptors, CXCR4 or CXCR7. Whether the observed changes are mediated by alterations in the activity of these receptors in hepatocytes or NPCs is unclear. Given the significant changes in *Ackr3* expression, specifically in hepatocytes, during fasting and refeeding, and the prominent alterations observed in mice consuming a WD, we sought to assess the impact of *Ackr3* overexpression in hepatocytes on hepatic lipid accumulation. To this end, 6-week-old mice were fed a WD for 2–3 weeks, similar to the *Cxcl12*-flox mice, and then injected via the tail vein with AAV*-Tbg::Ackr3* to induce hepatocyte-specific overexpression of *Ackr3* or AAV*-Tbg::GFP* as a control (Figure 3A, B).

**Figure 3.**
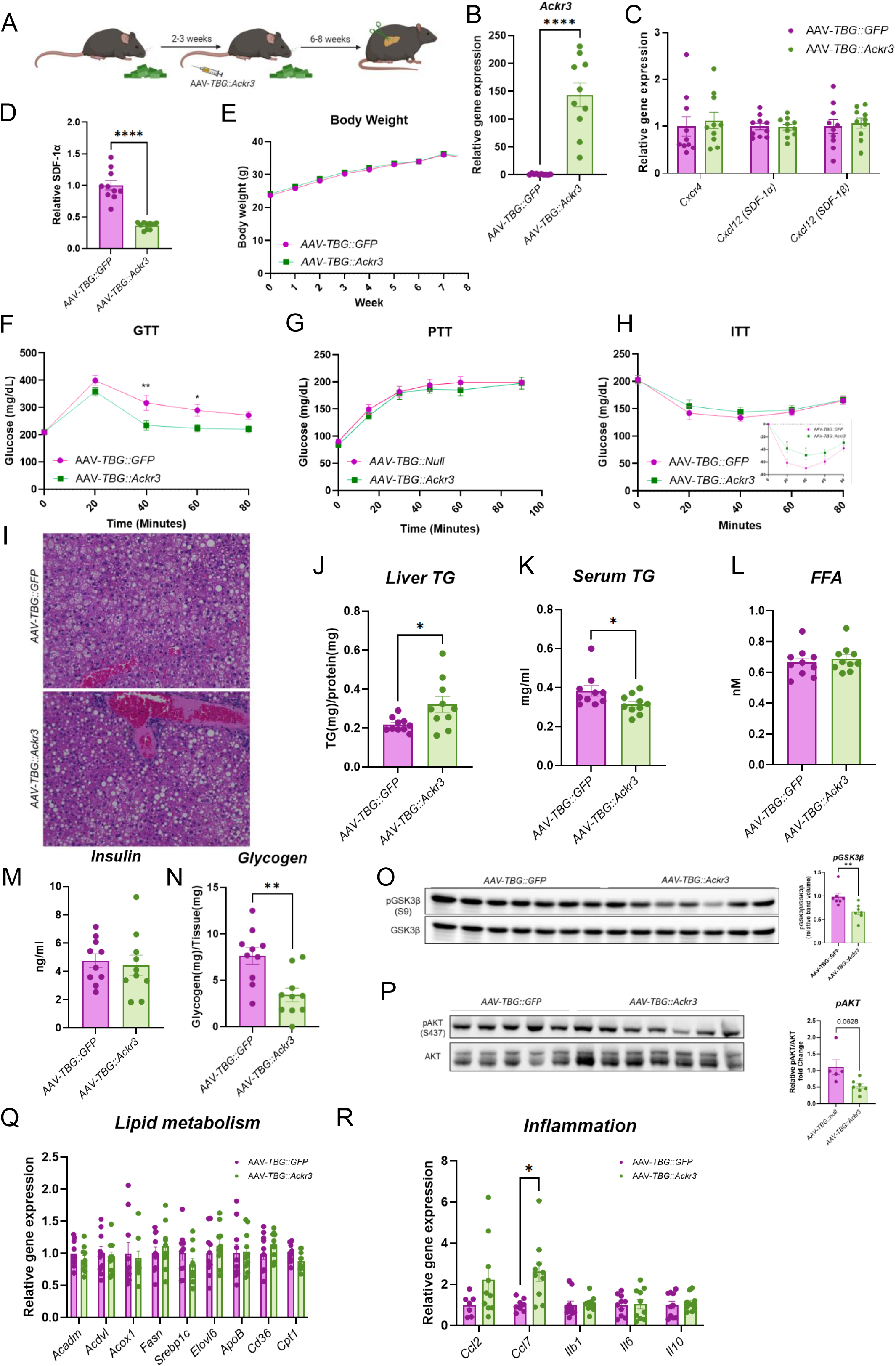
(**A**) Experimental design. *C57BL/6* mice were placed on a Western diet beginning at 6 weeks of age. Within each experiment, all mice were exposed to the diet for the same duration and analyzed together, although the exact duration varied slightly between independent cohorts. Mice were injected with AAV as indicated. Liver and serum samples were collected after 6hrs. of fasting. (**B**) *Ackr3* expression level in the livers of control and *AAV-TBG::Ackr3* mice. (**C**) *Cxcr4 and Cxcl12* expression levels in the liver. (**D**) SDF-1α levels in total liver lysates following SDF-1α ELISA. (**E**) Body weight. (**F, G, H**) Glucose, pyruvate and insulin tolerance tests. In the ITT the change from basal is in the insert. (**I**) H&E staining of liver sections from control and *Ackr3-*overexpression mice. (**J**) Quantification of liver TG (**K**), serum triglycerides (**L),** serum FFA (**M**), and serum Insulin. (**N**) Liver glycogen content. (**O**) pGSK3b (S9) in total liver extracts. (**P**) pAKT (S473) in total liver lysates (**Q**) Expression level of lipid metabolism-related and (**R**) inflammation-related genes. *, p-value<0.05; **, p-value<0.01; ****, p-value<0.0001.

Overexpression of *Ackr3* did not affect the endogenous expression levels of *Cxcr4* or *Cxcl12* (Figure 3C). Despite the absence of a change in hepatic *Cxcl12* mRNA expression, SDF-1α levels were markedly reduced in *Ackr3*-overexpressing livers (Figure 3D), consistent with enhanced ligand scavenging by CXCR7. Additionally, no differences in weight gain were observed between the groups (Figure 3E). An oral glucose tolerance test (GTT) revealed a modest reduction in the glucose excursion in mice overexpressing *Ackr3* (Figure 3F). In contrast, the pyruvate tolerance test (PTT) did not differ between groups (Figure 3G), suggesting that hepatocyte-specific *Ackr3* overexpression does not markedly alter pyruvate-driven glucose output under these conditions. Insulin tolerance testing showed only a subtle trend between groups (Figure 3H), while fasting insulin levels were unchanged (Figure 3M), indicating that the in vivo glucose phenotype is modest.

Importantly, similar to the effects observed with *Cxcl12* downregulation, *Ackr3* overexpression resulted in significant TG accumulation in the liver compared to the control group (Figure 3I, J). This was accompanied by a concurrent slight reduction in serum TG levels (Figure 3K), suggesting altered lipid metabolism. As seen in *Cxcl12*-downregulated mice, no significant differences were observed in serum FFA levels between the groups (Figure 3L). Interestingly, *Ackr3* overexpression resulted in reduced hepatic glycogen levels (Figure 3N). This reduction was accompanied by a decrease in the phosphorylation of GSK3β (Figure 3O), suggesting that impaired insulin signaling may be a potential contributing factor. Basal hepatic AKT phosphorylation following 6 hours of fasting showed a directional reduction in livers from *Ackr3*-overexpressing mice (Figure 3P), consistent with altered downstream AKT signaling in vivo. Despite the increase in hepatic triglyceride content, *Ackr3* overexpression did not significantly alter the expression of the lipid metabolism genes examined at this time point (Figure 3Q). Together with the reduction in hepatic glycogen and altered GSK3β phosphorylation, this suggests that the phenotype may reflect altered nutrient partitioning and signaling-dependent metabolic regulation rather than broad transcriptional remodeling of canonical lipid metabolism pathways. In contrast, significant upregulation of *Ccl7* and a slight, non-significant increase in *Ccl2* expression were observed in *Ackr3*-overexpressing livers, whereas the other inflammatory markers examined were unchanged (Figure 3R).

To further assess the metabolic phenotype of hepatocyte-specific *Ackr3* overexpression during the fasting-refeeding transition, we measured blood glucose in fasted and refed mice and analyzed livers collected in the refed state. Although differences in blood glucose were minor in the fasted state, *Ackr3*-overexpressing mice exhibited higher blood glucose after refeeding than control mice (Figure S4A). In the refed state, hepatic triglyceride content remained elevated, and glycogen content was reduced (Figure S4B,C). Because these biochemical measurements were obtained only after refeeding, they do not define the kinetics of fasting-refeeding adaptation, but they do indicate altered metabolic handling of the refed state in *Ackr3*-overexpressing mice. Refed-state gene-expression analysis revealed no broad changes in the lipid metabolism or inflammatory gene panels examined (Figure S4D,E), suggesting that this phenotype is not accompanied by major transcriptional remodeling at this time point.

These results show that hepatocyte-specific overexpression of *Ackr3* promotes hepatic TG accumulation and reduces glycogen stores in WD-fed mice, consistent with a role in hepatic energy metabolism and nutrient handling.

### Hepatocyte-specific overexpression of *Cxcr4* is not sufficient to modulate lipid and glucose metabolism

To further dissect the relative contribution of the CXCL12/CXCR4/CXCR7 axis to hepatic metabolism, we examined the effects of hepatocyte-specific overexpression of *Cxcr4*. CXCR7 is often described as a scavenger GPCR that internalizes and degrades its ligand, thereby modulating extracellular CXCL12 availability and indirectly influencing CXCR4 signaling(13). However, several reports have also suggested that CXCR7 can engage distinct intracellular signaling cascades, raising the possibility that the two receptors may exert both overlapping and independent functions in liver metabolism (13). It is therefore important to directly assess whether altering *Cxcr4* expression in hepatocytes is sufficient to recapitulate the effects observed with *Ackr3* manipulation.

Six-week-old mice were fed a WD for 2–3 weeks and subsequently injected via the tail vein with AAV-Tbg::*Cxcr4* to induce hepatocyte-specific overexpression of *Cxcr4* or AAV-*Tbg*::Null as a control (Figure 4A, B). Hepatocyte-specific *Cxcr4* overexpression resulted in a significant downregulation of *Ackr3* expression (Figure 4C), suggesting potential compensatory regulation within the axis. In contrast, hepatic SDF-1α levels were increased in the *Cxcr4* overexpression model (Figure 4D), suggesting that perturbation of receptor balance within the axis also influences ligand availability in vivo. Despite this, body weight gain was comparable between groups (Figure 4E). PTT revealed no differences in hepatic glucose output (Figure 4F), while ITT showed only a mild reduction in insulin sensitivity at 15 minutes post-insulin injection in the *Cxcr4*-overexpressing group (Figure 4G).

**Figure 4.**
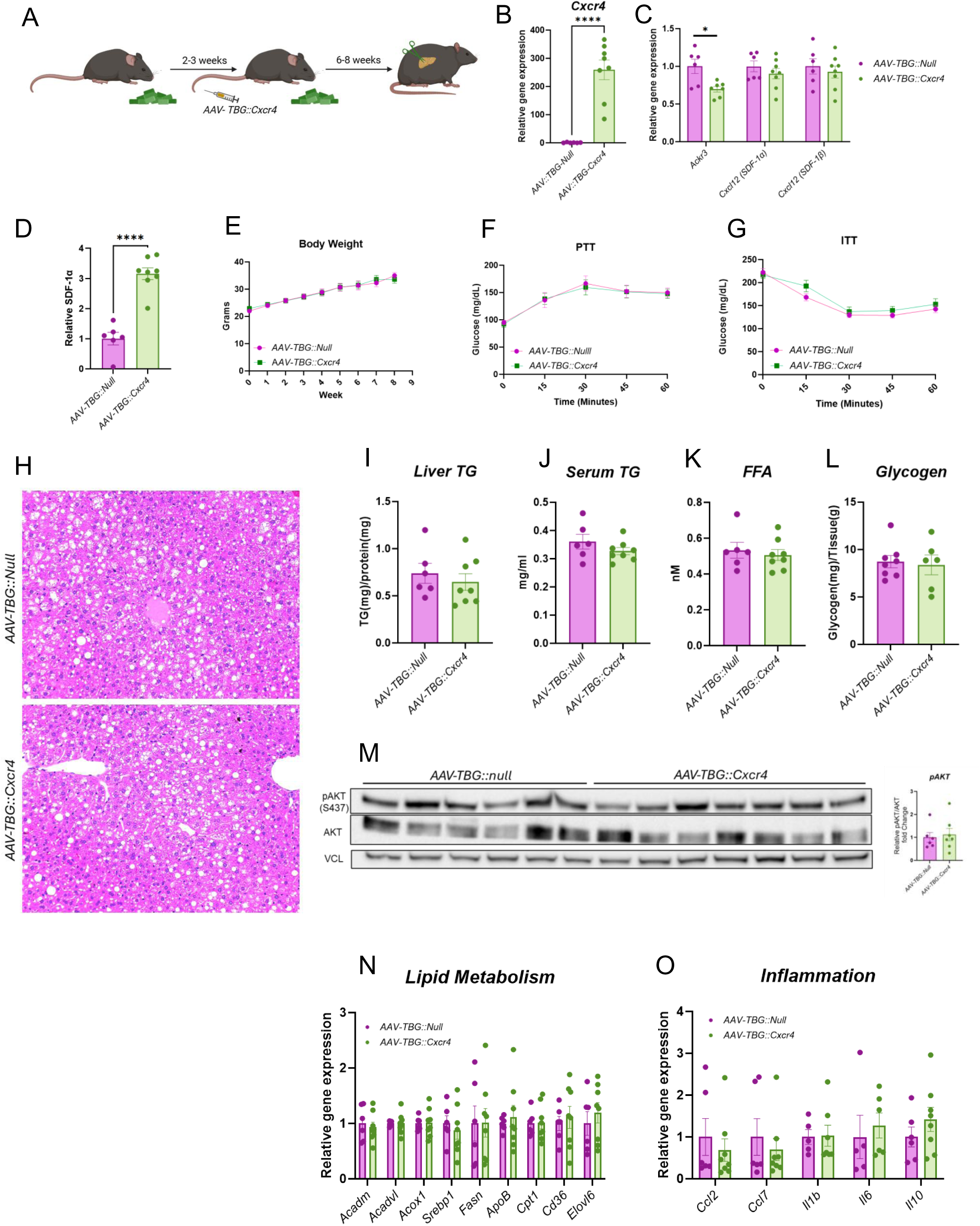
(**A**) Experimental design. *C57BL/6* mice were placed on a Western diet beginning at 6 weeks of age. Within each experiment, all mice were exposed to the diet for the same duration and analyzed together, although the exact duration varied slightly between independent cohorts. Mice were injected with AAV as indicated. Liver and serum samples were collected after 6hrs. of fasting. (**B**) *Cxcr4* expression level in the livers of *AAV-TBG::Cxcr4* and *AAV-TBG::Null* mice. (**C**) *Ackr3 and Cxcl12* expression levels in the liver. (**D**) SDF-1α levels in total liver lysates following SDF-1α ELISA. (**E**) Body weight. (**F, G**) Pyruvate and insulin tolerance tests. (**H**) H&E staining of liver sections from control and *Cxcr4-*overexpressing mice. (**I**) quantification of liver TG. (**J**), serum triglycerides (**K**), and serum FFA. (**L**) Liver glycogen content. (**M**) pAKT (S473) in total liver lysates (**N**) Expression level of lipid metabolism-related and (**O**) inflammation-related genes. *, p-value<0.05; ****, p-value<0.0001.

Histological analysis of H&E-stained liver sections showed no overt differences in morphology or lipid deposition (Figure 4H). Consistent with this, hepatic TG content, serum TG, and serum FFA levels were unchanged between groups (Figure 4I–K). Hepatic glycogen levels were also unaffected (Figure 4L). Basal hepatic AKT phosphorylation following 6 hours of fasting was not clearly altered by hepatocyte-specific *Cxcr4* overexpression (Figure 4M), indicating that increased *Cxcr4* expression alone is not sufficient to enhance downstream AKT signaling in this setting. Gene expression analysis of lipid metabolism pathways revealed no significant alterations in the expression of genes involved in β-oxidation, lipogenesis, or lipid handling (Figure 4N). Likewise, the expression of inflammatory markers, including *Ccl2* and *Ccl7*, was not altered (Figure 4O).

Taken together, these results indicate that hepatocyte-specific *Cxcr4* overexpression does not markedly affect hepatic lipid or glucose metabolism under WD feeding, in contrast to the changes observed with *Ackr3* overexpression. However, this model does not allow us to distinguish whether the effects of CXCL12 are mediated primarily through CXCR4, CXCR7, or a combination of both. Because *Cxcr4* overexpression also reduced *Ackr3* expression, receptor crosstalk remains a possible contributor. Definitive clarification will require complementary loss-of-function models to determine the relative contributions of CXCR4 and CXCR7 to hepatic metabolic regulation.

### Hepatocyte-specific CXCR7 overexpression enhances lipid accumulation and is associated with altered insulin signaling

Our results suggest that modulating *Cxcl12* and *Ackr3* expression in the liver is associated with lipid accumulation. To determine whether these effects can also be observed in a cell-autonomous manner, we injected mice with AAV-Tbg::*Ackr3* or AAV-Tbg::Null (control) via the tail vein and subsequently isolated primary hepatocytes. We then tested whether *Ackr3*-overexpressing hepatocytes display enhanced lipid accumulation in culture, consistent with the *in vivo* phenotype. Hepatocytes were incubated with an oleate/palmitate mixture (100 μM/50 μM) and stimulated with insulin (20 nM) to promote lipid storage. Neutral lipids were stained with Bodipy dye, visualized by fluorescence microscopy, and quantified by FACS analysis based on Bodipy intensity at the single-cell level. Our findings show that *Ackr3* overexpression markedly promotes lipid accumulation relative to controls, supporting the conclusion that the *in vivo* phenotype is at least partially mediated through cell-autonomous mechanisms (Fig. 5A) and indicating that *Ackr3* overexpression is sufficient to promote lipid accumulation in hepatocytes. To further examine the molecular changes associated with this phenotype, we analyzed insulin signaling in control and *Ackr3*-overexpressing hepatocytes. Across independent experiments, *Ackr3* overexpression was associated with a directional reduction in insulin-stimulated AKT phosphorylation, an effect that was more apparent in hepatocytes preincubated with the oleate/palmitate mixture (Figure 5B). Under these lipid-loading conditions, the effect of *Ackr3* on insulin signaling was therefore more evident than in untreated cells. Together, these data are consistent with altered insulin responsiveness in *Ackr3*-overexpressing hepatocytes, particularly in a lipid-rich context. We next asked whether mTORC1-linked signaling contributes functionally to the lipid accumulation phenotype. While *Ackr3* overexpression increased lipid accumulation, as described above, this effect was no longer observed when cells were treated with rapamycin (Figure 5C), indicating that the lipid-storage phenotype is sensitive to mTORC1 inhibition. These findings support the idea that mTORC1-linked signaling contributes to the pro-lipogenic effect of *Ackr3* in hepatocytes. It should be noted that these experiments were performed without exogenous CXCL12 stimulation. Primary hepatocytes secrete measurable and biologically active CXCL12 into the culture medium(29), providing an endogenous source of ligand. Because CXCR7 is known to sequester CXCL12 from the extracellular space, we considered the possibility that *Ackr3* overexpression alters hepatocyte signaling in part by changing ligand availability and thereby affecting CXCR4-dependent input. Consistent with this idea, hepatic SDF-1α levels were markedly reduced in the *Ackr3* overexpression model despite unchanged hepatic *Cxcl12* mRNA expression (Figure 3D), supporting the notion that increased CXCR7 expression can reduce ligand availability in vivo. In line with this interpretation, pharmacologic inhibition of CXCR4 with AMD3100 produced a similar directional reduction in insulin-stimulated AKT phosphorylation, particularly in lipid-loaded hepatocytes (Figure 5D). Together, the hepatocyte signaling experiments and the rapamycin-sensitive lipid phenotype, these findings support the idea that *Ackr3* overexpression is associated with altered insulin signaling that becomes more evident under lipid-rich conditions. In this context, our results suggest that increased CXCR7 expression under WD-feeding conditions may contribute to lipid accumulation and potentially promote the transition to steatohepatitis.

**Figure 5.**
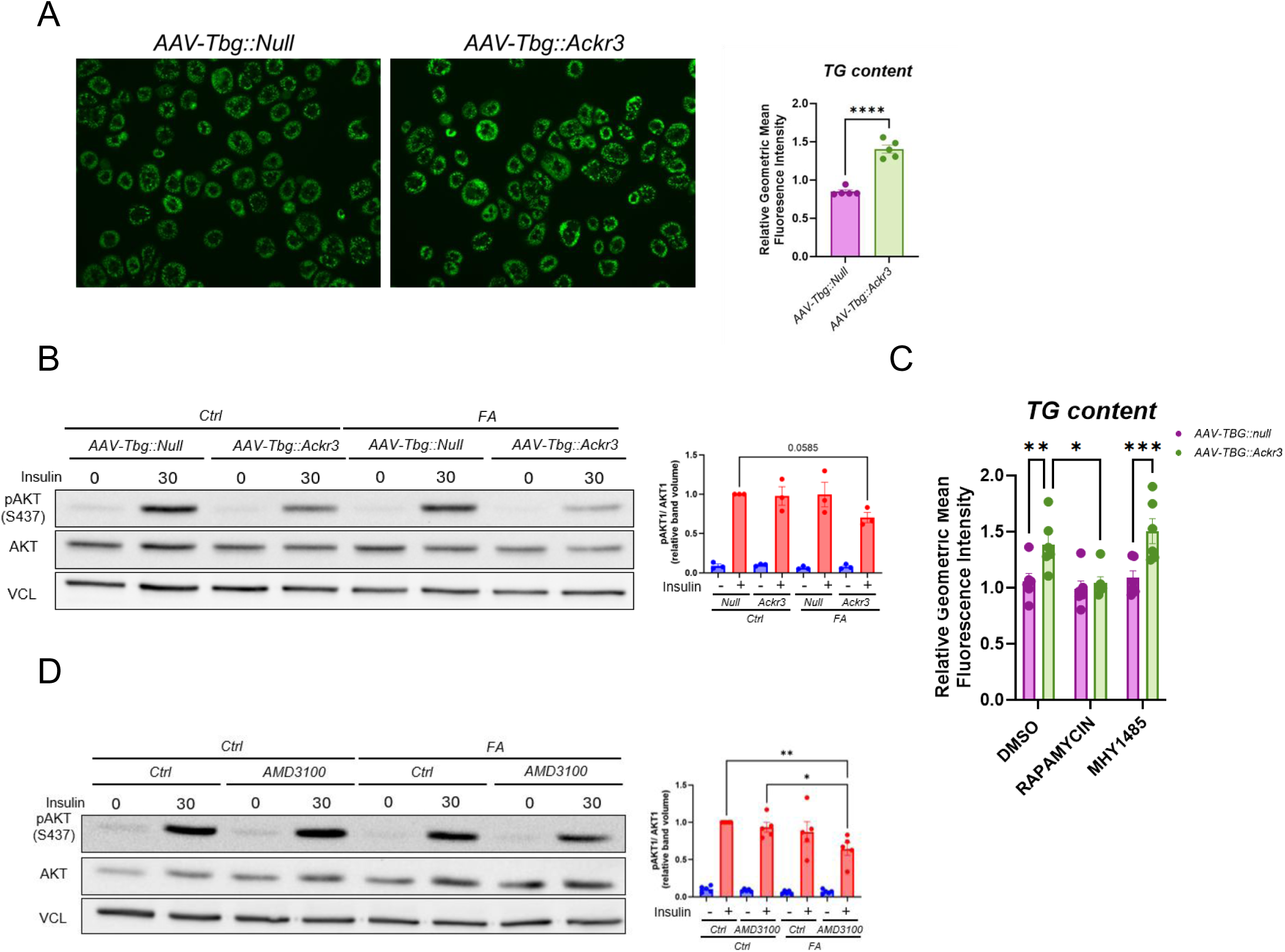
**(A)** Bodipy staining for lipid accumulation in control and *Ackr3*-overexpressing cultured primary hepatocytes, following a 20-hour treatment with oleic acid/palmitic acid mixture (100μM/50μM) and Insulin, 20nM (left panel). FACS analysis of Bodipy-stained treated cells (right panel). (**B**) Western blot analysis of control and *Ackr3*-overexpressing cultured primary hepatocytes, treated with 20 nM insulin for 0 and 30 minutes. FA, hepatocytes that were pretreated with oleic acid/palmitic acid mixture (100μM/50μM) for 20 hrs. before insulin stimulation. (**C**) FACS analysis of Bodipy-stained hepatocytes following loading with palmitic acid/oleic acid mixture and treatment with rapamycin (mTORC1 inhibitor) or MHY1485 (mTORC1 activator). (**D**) Western blot analysis of cultured primary hepatocytes treated with PBS or AMD3100 (CXCR4 antagonist) and with 20 nM insulin for 0 and 30 minutes. FA, hepatocytes that were pretreated with oleic acid/palmitic acid mixture (100μM/50μM) for 20 hrs. before insulin stimulation.

## Materials and Methods

### Animals

Wild-type (WT) C57BL/6OLAHsd mice were purchased from Envigo, and B6(FVB)-Cxcl12^tm1.1Link^/J (*Cxcl12-*flox) mice were obtained from The Jackson Laboratory (strain #021773). Only male mice were used in this study. Mice were housed under specific pathogen-free conditions with free access to food and water. The animal facility was maintained at 22°C (±2°C) with 40–70% relative humidity and a 12-hour light/dark cycle. The transgenic *in vivo* models mice were fed a high-fat/high sucrose Western diet (45% kcal fat/40% kcal carbohydrates, TD.08811, Teklad) starting at 6 weeks of age. After Western diet feeding, mice were allocated to the different AAV treatment groups based on body weight to achieve comparable baseline body weight distribution between groups. Sample sizes were based on prior experience with these in vivo models and endpoints. In most experiments, 6-8 mice were used per group. No animals were excluded except in cases of technical failure, such as unsuccessful injection, failed hepatocyte isolation, or inadequate sample quality. All experimental procedures were approved by the Hebrew University’s IACUC and conducted according to its guidelines.

### Primary mouse hepatocyte isolation and culture conditions

Primary hepatocytes were isolated using a Liberase perfusion technique. Eight-week-old WT mice were sacrificed by Isoflurane inhalation, and the liver was perfused via the vena cava with HBSS (0.5 mM EDTA, 25 mM HEPES, calcium-free, magnesium-free, phenol red-free; Sartorius Israel, Cat. 02-018-1A). During perfusion, the portal vein was clamped for 5 seconds after being cut. Subsequently, 10 mL of digestion buffer containing 25μg/ml Liberase was perfused(30). The liver was excised, rinsed with 10 mL of dPBS (Sartorius Israel, Cat. 02-023-1A), and gently scraped. The tissue was suspended in 20 mL of plating medium and passed through a 70-μm cell strainer. Live hepatocytes were separated using a Percoll gradient and plated in plating medium (high-glucose DMEM supplemented with 10% FBS, 2 mM pyruvate, 1% penicillin/streptomycin, 0.1 μM insulin, and 1 μM dexamethasone). After 3–4 hours, the medium was replaced with maintenance medium (low-glucose DMEM supplemented with 0.2% BSA, 4 mM L-glutamine, 1 mM pyruvate, 1% penicillin/streptomycin, 1 nM insulin, and 0.1 μM dexamethasone).

For hormonal stimulation experiments, hepatocytes were maintained in medium without hormones for 3 hours, followed by stimulation with either 20 nM insulin or 100 nM glucagon as indicated.

### Fasting and refeeding assays

Six-week-old WT mice were fed either a standard chow diet or a Western diet for 4 weeks. Hepatocytes were isolated after an ∼8-hour fasting period or after a 6-hour fast followed by 3 hours of refeeding. Hepatocytes were immediately separated from the NPCs and analysed for gene expression without plating. For *Cxcl12* gene expression analysis, total livers were collected from mice after sacrifice via Isoflurane inhalation and processed for further analysis.

### Plasmids, Adeno-Associated Viruses (AAVs), and transgenic models generation

The cDNA of *Ackr3* or *Cxcr4* was cloned into the pAAV.TBG.PI.eGFP.WPRE.bGH vector (Addgene #105535) using BamHI and NotI enzymes. AAV vectors AAV.TBG.PI.Cre.rBG (AAV-TBG::Cre) and pAAV.TBG.PI.Null.bGH (AAV-TBG::Null) were purchased from Addgene (catalog #107787 and #105536, respectively). AAVs were generated at the viral core facility of the Hebrew University in Jerusalem. To generate transgenic mice, a viral titer of 1.0 x10¹¹ genome copies in 100 μL saline was injected per mouse via the tail vein.

For *in vivo* hepatocyte-specific overexpression, WT C57BL/6 mice were injected with either *AAV-TBG::Ackr3* or *AAV-TBG::Cxcr4*, with control groups receiving *AAV-TBG::Null* or *AAV-TBG::GFP*. For hepatocyte-specific knockout of *Cxcl12*, *Cxcl12-*flox mice were injected with *AAV-TBG::Cre* or *AAV-TBG::Null* for the control group.

### Pyruvate, glucose, and insulin tolerance tests

Mice were fasted overnight (for the pyruvate tolerance test, PTT), 6 hours (for the glucose tolerance test, GTT), or 4 hours (for the insulin tolerance test, ITT). For PTT, mice were orally gavaged with a 25% pyruvate solution in Dulbecco’s PBS (dPBS) at a dose of 2.5 g/kg. For GTT, mice were gavaged with a 20% D-glucose solution in dPBS at a dose of 2 g/kg. For ITT, mice were intraperitoneally injected with 0.75 U/kg insulin. Blood glucose levels were measured at the indicated time points following injection.

### Serum free fatty acids, triglycerides, and insulin measurements

Mice were fasted for 6 hours before blood collection via cardiac puncture. The blood was allowed to clot, and serum was separated by centrifugation before being stored at –80°C for subsequent analysis. Serum free fatty acids (FFA) were measured using the NEFA-HR kit (FUJI FILM, Cat. 270-77000), triglyceride levels were assessed with the Triglyceride assay kit (Sigma-Aldrich, Cat. F6428 and T2449) in accordance with the manufacturer’s instructions, and serum insulin concentrations were determined using an ELISA kit (Alpco, Cat. CH01), following the manufacturer’s protocols.

### Hepatic triglyceride determination

Livers were collected from 6-hour-fasted mice, and triglycerides (TG) were extracted from approximately 50 mg of pulverized liver tissue. The tissue was homogenized in 800 µL of PBS, and 400 µL of the homogenate was mixed with 800 µL of a 2:1 chloroform:methanol solution. After centrifugation at maximum speed, the lower lipid phase was transferred to a new tube and evaporated overnight in a chemical hood. The resulting lipids were reconstituted in 50 µL of a 60% butanol and 40% Triton-114:methanol (2:1) solution. TG concentrations were measured after a 1:4 dilution with ethanol using the Triglyceride assay kit. Results were normalized to protein content, determined by the QPro-BCA assay (Cyanagen).

### Liver glycogen determination

Approximately 50 mg of pulverized liver tissue was homogenized in 0.6 N perchloric acid (5 volumes per tissue weight). The homogenate was centrifuged at 13,000 × g for 10 minutes, and 80 µL of the supernatant was neutralized with 40 µL of 1 M KHCO_₃_. The neutralized sample was diluted 1:20 in 0.4 M acetate buffer (pH 4.8) (8.09g Sodium Acetate trihydrate, 2.4ml 99% acetic acid, and made up to 500mL in distilled deionized water).

For glycogen quantification, 10 µL of the diluted sample or glycogen standard was incubated with 50 µL of 1 mg/mL amyloglucosidase (Sigma-Aldrich, Cat. A7420), prepared in acetate buffer, for 2 hours at 37°C. Basal glucose levels were determined by incubating the diluted samples with acetate buffer alone. After incubation, glucose levels were measured using the Glucose (GO) Assay Kit (Sigma-Aldrich, Cat. GAGO20), following the manufacturer’s protocol. Glycogen content was calculated by subtracting the basal glucose level from the total glucose level and comparing the results to a standard curve.

### Liver SDF-1α determination

SDF-1α content was measured using a commercial ELISA kit (Mouse CXCL12/SDF-1 alpha Quantikine ELISA Kit, R&D, Cat# MCX120). ∼100mg of pulverized liver tissue were homogenized in 500ml dPBS supplemented with protease inhibitors cocktail. The homogenate was cleared by centrifugation (10min, X10,000g) and 50μl of cleared homogenate were used directly for the assay which was carried out according to the manufacturer’s instructions.

### Real-time quantitative PCR analysis

Total RNA was extracted from primary hepatocytes or liver tissues using the NucleoSpin® RNA kit (Takara Bio). One microgram of RNA was reverse-transcribed into cDNA using the Qscript™ cDNA Synthesis Kit (QuantaBio, Cat. #95047-500). Quantitative PCR (qPCR) was performed using the Luna Universal qPCR Master Mix (NEB, Cat. M3003E) on a C1000 Touch Thermal Cycler equipped with the CFX384 system (Bio-Rad). Gene expression levels were normalized to the housekeeping genes β*2M* and *Rpl13*. Data analysis was conducted using the Bio-Rad CFX Maestro software.

### Hematoxylin and Eosin staining

Liver tissues were collected at the end of *in vivo* experiments, fixed in 10% formalin for 24 hours, and then stored in 70% ethanol. Hematoxylin and eosin (H&E) staining was performed following a standard protocol. Images were captured using an Evos M5000 microscope and 20× magnification.

### Immunohistochemistry

The following primary antibodies were used: anti-F4/80 (Bio-Rad, cat. MCA497G, clone CI:A3-1) at 1:200 dilution with Proteinase K antigen retrieval; anti-CD19 (Thermo Fisher, cat. 14-0194-82, clone 6OMP31) at 1:500 dilution following citrate antigen retrieval (pH 6.0); and anti-CD3 (Bio-Rad, cat. MCA1477, clone CD3–12) at 1:200 dilution with EDTA-based antigen retrieval (pH 8.0). Immunohistochemistry was performed on formalin-fixed paraffin-embedded (FFPE) sections. Sections were de-paraffinized and rehydrated, followed by 2 washes in DW. Antigen retrieval was done in Citrate or Tris buffers in a pressure cooker (125Lc, 3 min) for CD3 and CD19, or by incubation with Proteinase K for F4/80. Sections were rinsed twice in DW and incubated in 3% Hydrogen Peroxide for 10 min at RT, followed by 3 rinses in DW. Sections were washed in wash buffer and blocked with protein block for 2 hours at RT. Sections were incubated with primary antibody diluted in protein block overnight at 4Lc and washed 3 times in wash buffer. Sections were incubated with HRP-conjugated secondary antibodies for 30 min at RT and washed in wash buffer. DAB solution was added for up to 15 min in the dark, and sections were rinsed in tap water. Hematoxylin staining was done for 10 to 60 seconds, followed by washes in tap water. Sections were dehydrated and mounted.

### Bodipy staining

Three hours after primary hepatocytes were isolated, the plating medium was replaced with maintenance medium supplemented with 100 µM oleic acid (Sigma-Aldrich, Cat. O1383) and 50 µM palmitic acid mixture for 20 hours. Following incubation, the medium was removed, and the cells were washed with dPBS. Cells were then incubated with Bodipy dye (1:2500 dilution of 5mM stock in dPBS) for 15 minutes at 37°C. After staining, the cells were washed twice with dPBS and fixed with 4% paraformaldehyde for 30 minutes at room temperature. Fixed cells were washed again twice with dPBS, and images were captured using an Evos M5000 microscope (20× magnification). For flow cytometry, Bodipy-stained cells were trypsinized in the presence of DNase I for 1 minute at 37°C, neutralized by the addition of fetal bovine serum (FBS), collected and centrifuged, and fixed using Cytofix/Cytoperm (BD Biosciences) for 20 minutes at room temperature. Cells were then centrifuged, resuspended in Perm/Wash buffer (BD Biosciences), and analyzed by flow cytometry. Data were analyzed using FlowJo software, and geometric mean fluorescence intensity (gMFI) was calculated(31).

### Statistical Analysis

Data are presented as mean ± SEM. Statistical comparisons were performed using GraphPad Prism 9.0. ANOVA with Sidak post hoc analysis or an unpaired two-tailed Student’s t-test was used. A *p*-value of <0.05 was considered statistically significant.

## Discussion

In this study, we identify a previously unrecognized role for the CXCL12/CXCR4/CXCR7 axis in hepatocyte metabolic regulation. Beyond prior work linking this pathway to liver injury, fibrosis, regeneration, and cancer(7, 10, 16), our findings show that this chemokine network is dynamically regulated during the fasting-refeeding transition and contributes directly to hepatic lipid homeostasis. By combining hepatocyte-specific manipulation of *Cxcl12*, *Ackr3*, and *Cxcr4*, we show that reduced hepatocyte-derived CXCL12 and increased hepatocyte CXCR7 promote hepatic triglyceride accumulation, whereas hepatocyte *Cxcr4* overexpression does not phenocopy this response. These findings support a nonredundant, hepatocyte-autonomous role for CXCR7 and connect this chemokine axis to insulin/mTORC1-linked metabolic regulation and lipid storage.

Our data reveal that *Ackr3* expression is induced by glucagon and suppressed by refeeding, while *Cxcr4* is downregulated by insulin. A similar reduction in *Cxcr4* expression following insulin stimulation has also been described in endothelial cells(25). This reciprocal pattern links the CXCL12/CXCR4/CXCR7 axis to the opposing hormonal cues that orchestrate hepatic adaptation to nutrient status. Under Western-diet conditions, however, this regulation becomes disrupted: *Ackr3* expression remains aberrantly elevated during refeeding, and *Cxcr4* repression is blunted. These alterations suggest that chronic nutrient excess impairs chemokine-receptor homeostasis, potentially contributing to the dysregulated lipid handling and insulin signaling that characterize steatotic livers. The increase in hepatic SDF-1α observed in the *Cxcr4* overexpression model is also notable (Figure 4D). One possible explanation is that elevated CXCR4 expression shifts receptor balance within the axis in a manner that suppresses *Ackr3* expression, as observed in our data, thereby reducing CXCR7-dependent ligand scavenging and allowing hepatic SDF-1α to accumulate. More broadly, this result suggests that receptor abundance within the CXCL12/CXCR4/CXCR7 system may feed back on ligand availability in vivo, further supporting the idea that this pathway operates as an integrated regulatory axis rather than as independent ligand-receptor pairs.

Our findings reveal that CXCR7 functions as a hepatocyte-autonomous regulator of lipid accumulation and insulin responsiveness, acting independently of systemic or paracrine influences. Hepatocyte-specific overexpression of *Ackr3* was sufficient to induce steatosis *in vivo* and to promote lipid droplet formation in isolated hepatocytes maintained without exogenous CXCL12, indicating that the effect may originate, at least in part, within hepatocytes. Notably, hepatic triglyceride accumulation in the *Ackr3* overexpression model was not accompanied by broad changes in the lipid metabolism gene panel examined, suggesting that the phenotype may arise primarily from signaling-dependent changes in nutrient partitioning and lipid storage rather than from large transcriptional shifts in canonical lipogenic or oxidative pathways. Mechanistically, CXCR7 behaves as an atypical β-arrestin–biased receptor that does not engage classical G-protein signaling but instead signals through β-arrestin scaffolds(32). Several studies have shown that β-arrestin signaling significantly affects mTOR by regulating the upstream Akt and ERK1/2 signaling pathways, which subsequently converge to activate mTOR Complex 1 (mTORC1) and promote protein translation(33–35). This raises the possibility that CXCR7-mediated β-arrestin signaling modulates mTORC1-linked signaling in hepatocytes, thereby influencing downstream metabolic responses linked to lipid storage. Consistent with this model, *Ackr3*-overexpressing hepatocytes displayed increased lipid droplet formation, which was abolished by rapamycin, supporting involvement of mTORC1-linked signaling in the lipogenic phenotype. This β-arrestin-mTOR interaction may enable CXCR7 to promote a more anabolic metabolic state in hepatocytes. In parallel, CXCR7 functions as a high-affinity scavenger receptor that internalizes and degrades CXCL12(36, 37), thereby reducing ligand availability for CXCR4 and attenuating CXCR4-mediated PI3K-Akt signaling that normally promotes insulin sensitivity and suppresses gluconeogenesis(11, 13, 22). In line with this possibility, hepatic SDF-1α levels were markedly reduced in the *Ackr3* overexpression model despite unchanged hepatic *Cxcl12* mRNA expression, supporting the idea that increased CXCR7 expression can alter ligand availability in vivo. The combined outcome may be a signaling imbalance characterized by reduced CXCR4 input, altered insulin responsiveness, and enhanced lipid storage. The in vivo pAKT analyses across the three models were also consistent with this interpretation, as reduced ligand availability in the *Cxcl12* loss-of-function model and increased scavenging in the *Ackr3* overexpression model were each associated with lower hepatic AKT phosphorylation, whereas *Cxcr4* overexpression alone did not increase this signal. Under lipid-rich conditions, this imbalance may become more apparent, consistent with the reduced insulin-stimulated AKT phosphorylation observed in hepatocytes. Importantly, in the present study, these findings are interpreted in the context of hepatocyte metabolic signaling and lipid handling, as primary hepatocytes in our experimental setting are non-proliferative, and we did not directly assess survival or apoptosis endpoints. Together, these findings position CXCR7 as a cell-autonomous integrator of chemokine and nutrient-sensing pathways in hepatocytes.

Although both CXCR4 and CXCR7 receptors bind CXCL12, their signaling properties are fundamentally distinct yet intricately coordinated. CXCR4 is a canonical GPCR that couples to Gα_i_ proteins, activating PI3K–Akt, ERK, and calcium signaling to promote insulin sensitivity, cell survival, and metabolic adaptation(12). In contrast, CXCR7 lacks efficient G-protein coupling. Instead, it functions as an atypical, β-arrestin–biased receptor that signals through arrestin-dependent scaffolds while simultaneously scavenging and internalizing CXCL12 to regulate its extracellular availability, as discussed above(32, 38). Recent structural and mechanistic work has revealed that these receptors communicate through phosphorylation-dependent crosstalk(14, 15). These studies show that CXCR7 can “sense” CXCR4 activation via GRK-mediated phosphorylation barcodes on CXCR7, which serve as docking cues for β-arrestin binding(15). They further demonstrated that distinct CXCR7 phosphorylation patterns modulate the strength and duration of β-arrestin recruitment to CXCR7, thereby tuning the amplitude of CXCR7 signaling(14). Through this cross-receptor sensing, CXCR7 interprets the activation state of CXCR4 and converts it into β-arrestin–dependent outputs, effectively linking ligand-induced CXCR4 activity to downstream CXCR7 signaling. In hepatocytes, such coupling likely determines how CXCL12 gradients are translated into metabolic responses. Overexpression of CXCR7 may therefore enhance ligand scavenging, limiting CXCR4-driven Akt activation while also shifting mTORC1-linked metabolic signaling, actions consistent with our observations of reduced SDF-1α availability, altered insulin responsiveness, and rapamycin-sensitive lipid accumulation. Conversely, CXCR4 overexpression suppressed *Ackr3* expression, suggesting a compensatory mechanism to maintain receptor balance. Consistent with this idea, hepatic SDF-1α levels were increased in the *Cxcr4* overexpression model, possibly reflecting reduced CXCR7-dependent ligand scavenging. Together, these findings indicate that hepatic metabolism is governed not by isolated receptor activities but by a dynamic CXCR4-CXCR7 partnership in which GRK-dependent phosphorylation, β-arrestin recruitment, and CXCL12 scavenging collectively shape the balance between anabolic and insulin-sensitizing pathways in hepatocytes. Mechanistically, our data support a model in which CXCL12 availability and CXCR7 expression together influence hepatocyte insulin signaling and lipid storage, implicating CXCR7 alongside CXCR4 in modulating hepatic energy metabolism.

An additional point to consider is ligand specificity within this pathway. Although CXCR7 can bind ligands beyond CXCL12, including CXCL11, the present study was performed in C57BL/6 mice, in which *Cxcl11* is not functionally expressed due to a naturally occurring inactivating mutation (39). CXCL11 is therefore unlikely to make a substantial contribution to endogenous hepatic CXCR7 signaling in our models. This issue may, however, be more relevant in human liver biology, where CXCR7 can engage a broader ligand repertoire. Likewise, because our study relied primarily on hepatocyte-specific genetic and gain-of-function approaches, future studies using pharmacologic CXCR7 inhibition and complementary loss-of-function strategies will be important for further resolving the relative contributions of ligand scavenging and downstream receptor signaling to hepatic metabolic regulation.

Together, our findings position the CXCL12/CXCR4/CXCR7 axis as an integrative signaling hub that links chemokine communication with hepatic energy metabolism. Under physiological conditions, CXCL12 derived from hepatocytes and non-parenchymal cells coordinates CXCR4 and CXCR7 activity to maintain lipid balance, insulin responsiveness, and adaptation to fasting and feeding. Glucagon and insulin fine-tune this axis by reciprocally regulating *Ackr3* and *Cxcr4* expression, ensuring that chemokine signaling aligns with the prevailing hormonal and nutritional state. Under metabolic stress, however, this coordination is impaired: excessive nutrient load or insulin resistance is associated with aberrant CXCR7 activation and loss of CXCR4 input, creating a signaling bias that favors altered mTORC1 responsiveness, triglyceride accumulation, and inflammatory gene expression. The altered expression patterns of *CXCL12*, *CXCR4*, and *CXCR7* observed in human NAFLD and NASH datasets support the possible relevance of this pathway to metabolic liver disease. These findings suggest that CXCR7 may be relevant to restoring hepatic metabolic balance. Modulating CXCR7 activity, either by limiting its ligand-scavenging capacity, rebalancing CXCL12 availability, or selectively tuning its β-arrestin bias, may influence CXCR4-Akt signaling and mTORC1-linked metabolic regulation in hepatocytes. Furthermore, elucidating how GRK-mediated phosphorylation and β-arrestin recruitment operate in hepatocytes across different nutritional and hormonal contexts will be important for a better understanding of how chemokine receptor balance shapes hepatic metabolism. Collectively, our work extends the conceptual framework of the CXCL12/CXCR4/CXCR7 axis beyond its canonical roles in fibrosis, angiogenesis, and cancer to a new function in hepatic metabolic regulation and may warrant future evaluation as a candidate therapeutic axis in metabolic liver disease.

## Acknowledgments

This work was supported by the Israel Science Foundation (ISF grant 1280/21) to KS. We thank all members of the laboratory for their valuable discussions and technical assistance throughout the study.

## Author contributions

R.B.M. conceptualized, conducted, analyzed experiments and prepared data for publication. C.A.T, H.B, N.H, E.C and I.S, conducted experiments. K.S, conceived the project, conceptualized, conducted experiments, wrote the manuscript, and prepared data for publication.

## Declaration of generative AI and AI-assisted technologies in the manuscript preparation process

During the preparation of this work, the authors used ChatGPT in order to edit and improve the readability of the text. After using this tool/service, the authors reviewed and edited the content as needed and take full responsibility for the content of the published article.

---

**Figure S1.**
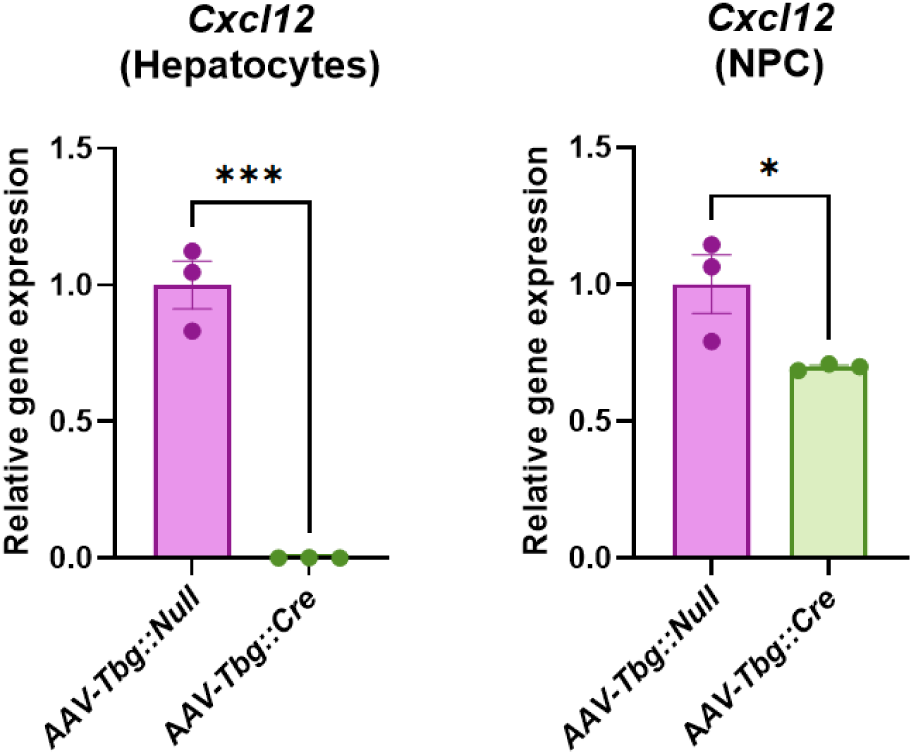
*Cxcl12*-flox mice were injected with AAV8-*TBG*-Cre, and livers were subjected to collagenase perfusion as detailed in the Methods. Primary hepatocytes were separated from non-parenchymal cells (NPCs), and total RNA was immediately extracted for analysis. *Cxcl12* expression was completely ablated in hepatocytes, whereas its expression in NPCs was only partially reduced.*, p-value<0.05; ***, p-value<0.001.

**Figure S2.**
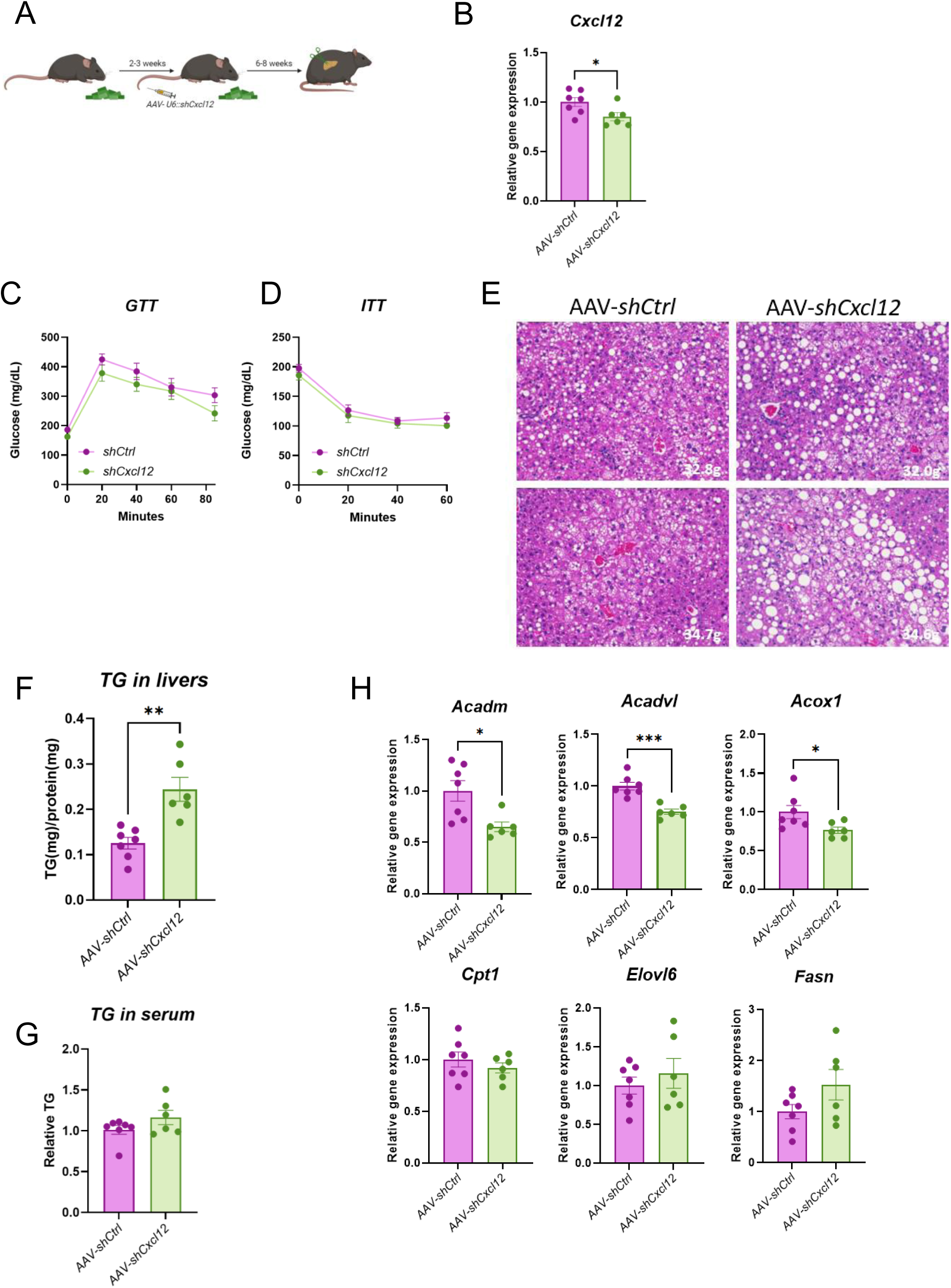
(**A**) Experimental design. *C57BL/6* mice were placed on a Western diet beginning at 6 weeks of age. Within each experiment, all mice were exposed to the diet for the same duration and analyzed together, although the exact duration varied slightly between independent cohorts. Mice were injected with AAV as indicated. (**B**) *Cxcl12* expression level in the liver following delivery of AAV-shCtrl or AAV-sh*Cxcl12.* (**C**) Glucose and (**D**) insulin tolerance tests. (**E**) H&E staining of liver sections from *Cxcl12*-depleted mice. (**F, G**) Quantification of liver and serum TG (**H**) expression level of lipid metabolism-related genes. *, p-value<0.05; **, p-value<0.01; ***, p-value<0.001.

**Figure S3.**
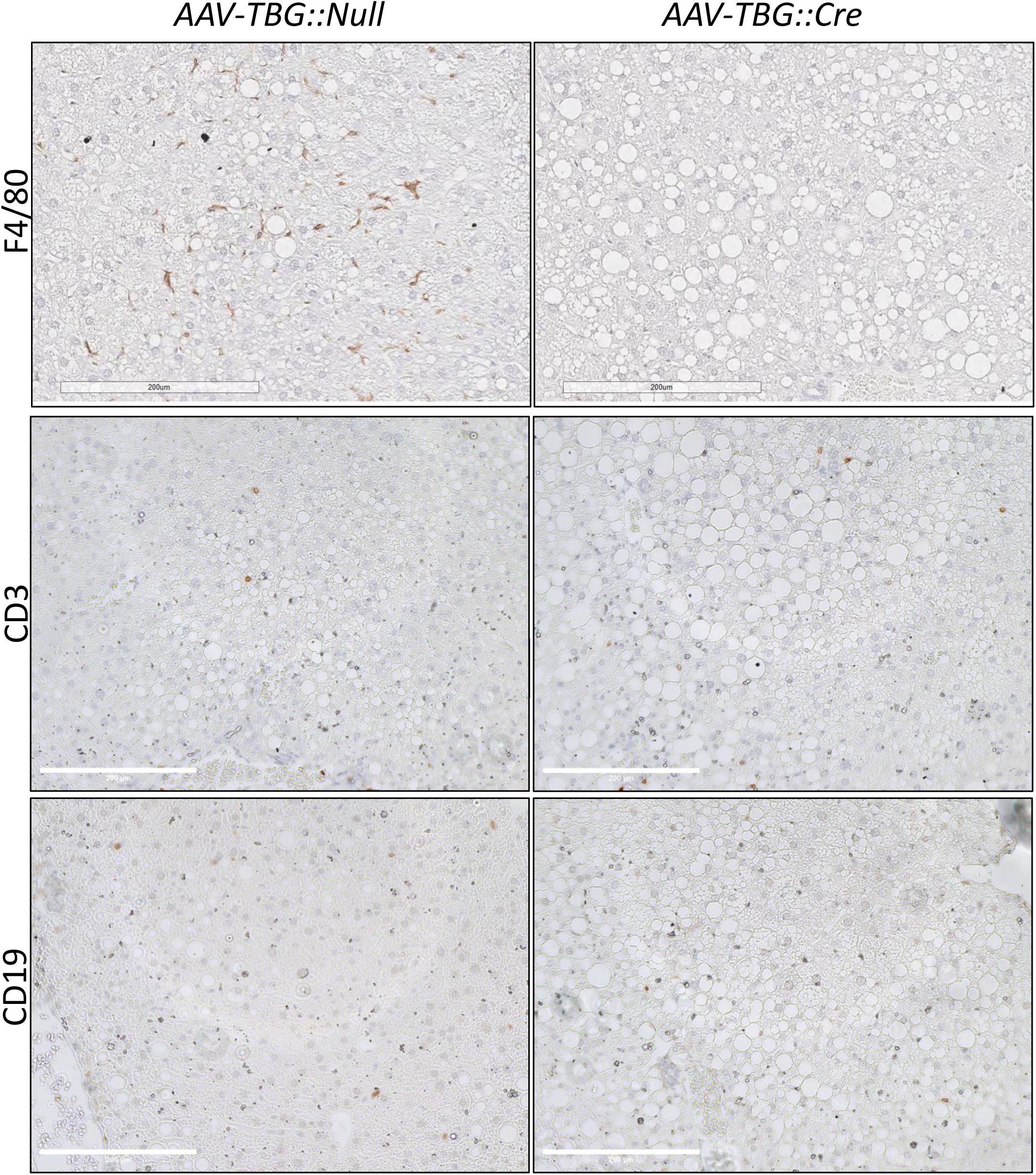
Immunostaining for macrophages (F4/80), T cells (CD3), and B cells (CD19) in liver sections from control (AAV-*TBG::Null*) and *Cxcl12*-depleted (AAV-*TBG*::*Cre*) mice.

**Figure S4.**
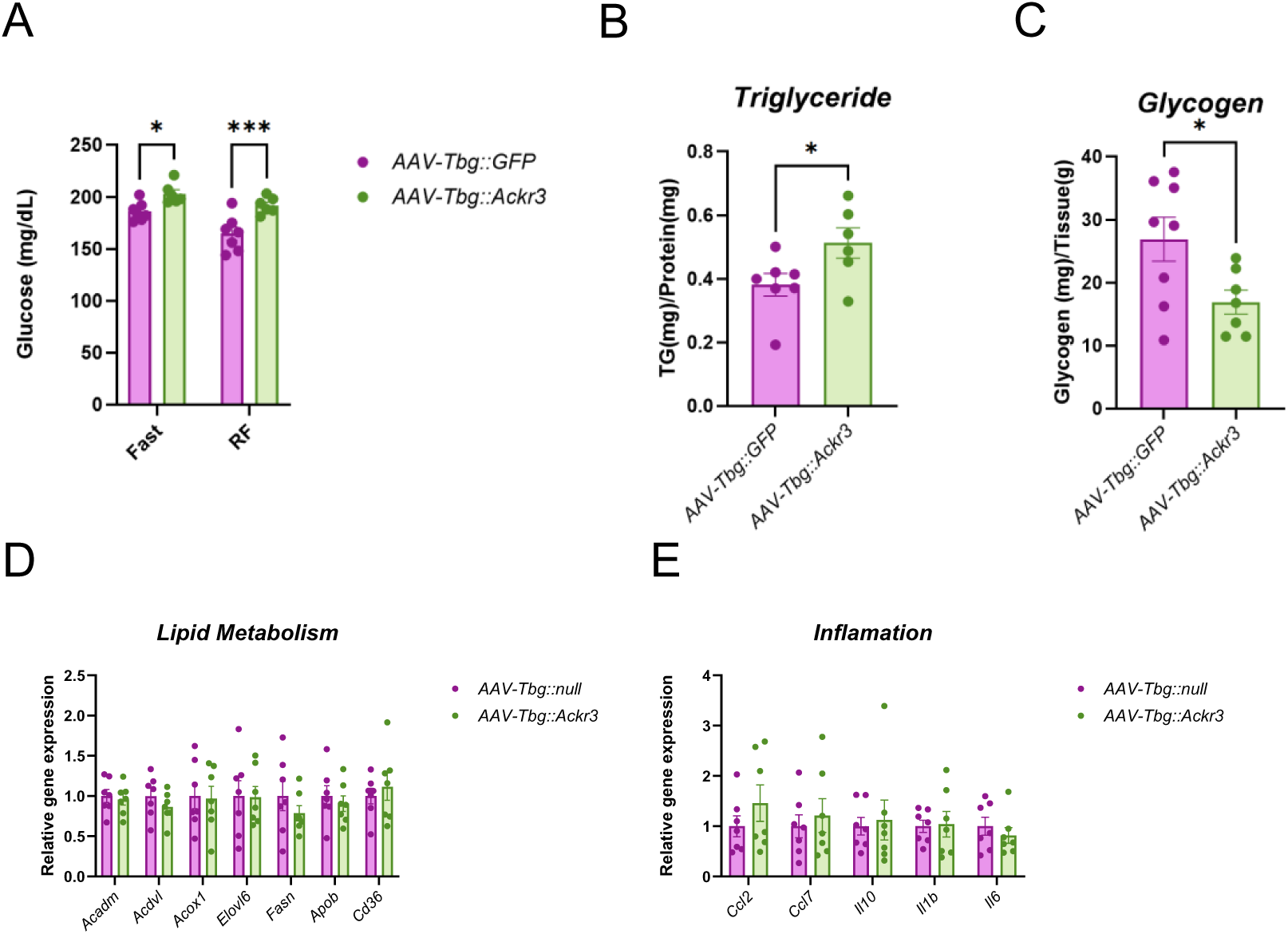
(**A**) Blood glucose concentrations in *Ackr3*-overexpressing mice following fasting (6 hrs.) and refeeding (RF, 1 hr.). (**B**) Liver TG and (**C**) liver glycogen levels in *Ackr3*-overexpressing mice. The liver tissues were collected in the refed state following 6 hrs. of fast and then 3 hrs. of refeeding. Expression levels of lipid metabolism (**D**) and inflammatory genes (**E**) from liver tissues collected in the refed state. *, p-value<0.05; ***, p-value<0.001.

